# Peristaltic contractions drive gut anisotropic growth through collective cell rearrangements

**DOI:** 10.64898/2025.12.22.695864

**Authors:** Koji Kawamura, Yoshiko Takahashi, Masafumi Inaba

**Affiliations:** Department of Zoology, Graduate School of Science, Kyoto University Kitashirakawa, Sakyo-ku, Kyoto, Japan

**Keywords:** Gut elongation, peristalsis, chicken embryo, optogenetics, live-cell imaging

## Abstract

The gut is the longest organ in the body, and its length is essential for efficient absorption of nutrient in adults. It is known that the gut undergoes massive elongation during prenatal/embryonic stages^1,2^, in which anisotropic growth takes place. But how this characteristic organ growth is achieved remains unknown. We here demonstrate an unprecedented role of peristaltic movements in the embryonic gut elongation using a caecum as a novel model in chicken embryos. Inhibition of the peristaltic movements by chemical drugs impedes the elongation of caecum. The elongation-halted caecum restores its longitudinal growth by optogenetically introduced peristaltic movements. At the cellular level, we have found that circular smooth muscle cells divide along the circumferential axis in a peristalsis-independent manner, which would otherwise serve as an engine for the radial growth. However, with peristaltic stimulation, the cells expand collectively their distribution along the gut longitudinal axis leading to the elongation of this organ. Thus, peristaltic activity biases circumferentially proliferating smooth muscle cells toward longitudinal rearrangement. This two-step model offers a novel way to identify the peristaltic activity as a critical growth signal for the gut elongation, and also to parsimoniously explain the anisotropic growth of the gut, which undergoes the elongation while retaining its thickness.

## Introduction

The gut is the longest organ in the body, and its length is essential for efficient absorption of nutrient. In adult humans, the small intestine reaches ∼5.5 m, and much of this length is achieved during prenatal/embryonic growth ^1,2^. What enables this massive elongation remains poorly explored. For the gut tube elongation, anisotropic growth is required in which the longitudinal growth must predominate over the radial growth. Although previous studies in several model organisms reported oriented cell divisions and intercalations of endodermal epithelial cells that occur locally in the gut, it remains unclear whether these events match the organ level scale of massive elongation ^3–5^.

In the course of studies we have been conducting with embryonic gut in chickens, we witnessed a massive elongation of this organ. The entire gut tube (posterior to stomach) increases its length by 1.91-fold in two days from embryonic day 10 (E10) to E12, and this growth rate is overwhelming over other organs; the body (1.24), forelimb (1.40), and eye (1.10) (Fig. 1a, b) ^6^. The stages E10 to E12 coincide with the onset of prominent movement of this organ, called gut peristalsis, delivering periodic and progressive waves of local contraction ^7–9^. However, due to the convoluted structures, the main tract (midgut and hindgut) hampers elaborate scrutinization that requires accurate measurement and quantitative analyses.

**Figure 1:**
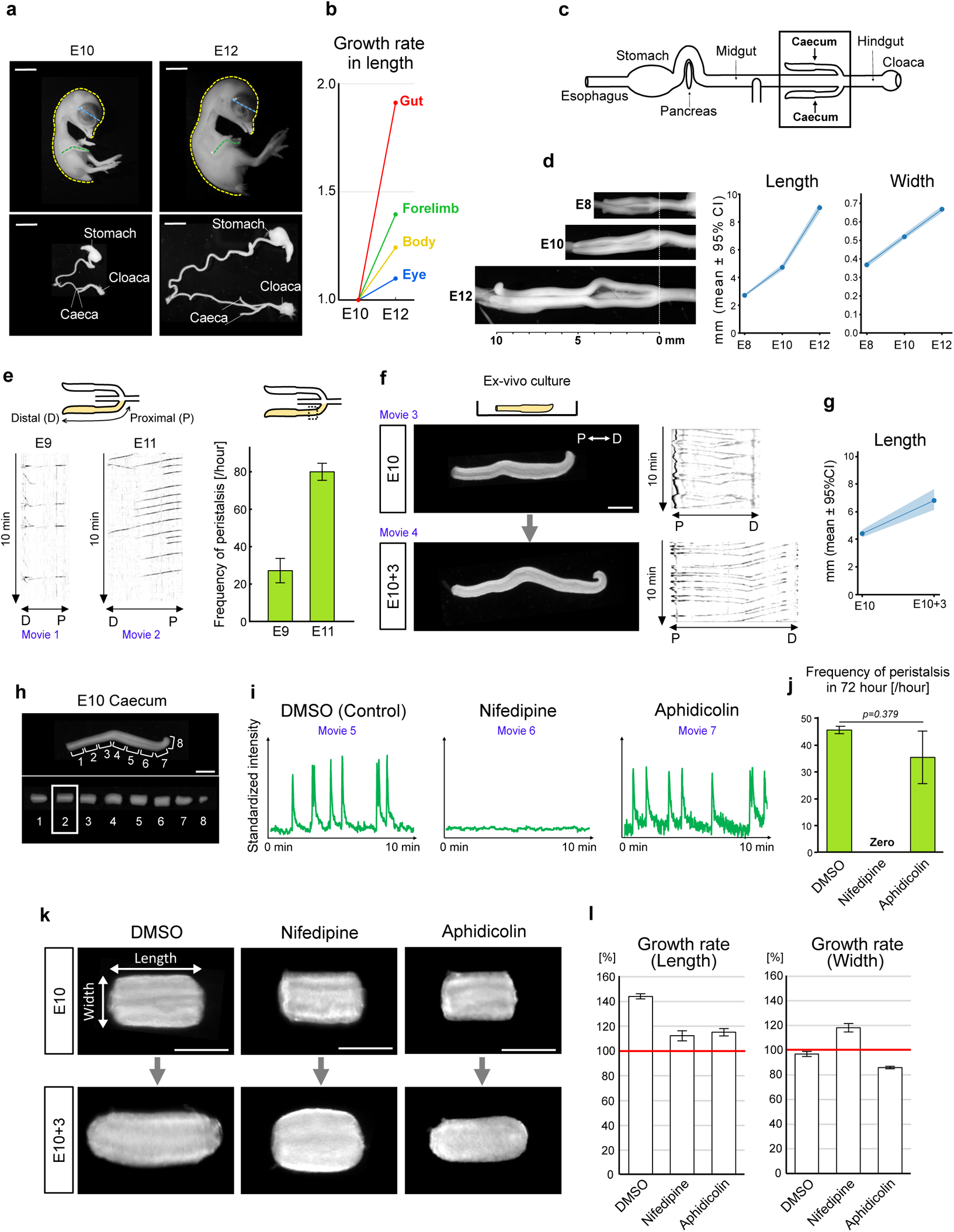
Gut peristalsis and cell proliferation are required for caecum elongation. **a**, Representative images of whole chicken embryos (upper panels) and digestive tracts (lower panels) at embryonic day 10 (E10) and E12. Scale bars, 10 mm. **b**, Growth of the gut, forelimb, body, and eye between E10 and E12. Each point represents the mean value from three embryos. **c**, Schematic illustration of the chicken digestive tract. **d**, Developmental changes in a pair of caeca from E8 to E12 (left). Graphs (right) show changes in caecal length and width (mean ± 95% CI) at E8 (n = 16), E10 (n = 24), and E12 (n = 20). **e,** Kymographs of peristaltic waves in E9 (left) and E11 (right) caeca. The horizontal axis and vertical axis represent position along the caecal axis and time, respectively. Bar graphs (right) show the frequencies of peristaltic waves passing through the proximal region of the caecum indicated by the dotted area in the schematic. **f**, Ex-vivo culture of the caecum. Left column: representative images of caeca at E10 (upper) and after 3 days in culture (lower); a schematic of the culture system is shown at the top. Right column: kymographs corresponding to each condition. Proximal (P)-distal (D) axes are indicated in both images and kymographs. Scale bar, 1 mm. **g**, Changes in caecal length during ex vivo culture (mean ± 95% CI; n = 6). **h**, Representative images of the caecum showing the whole view (upper) and a fragmented view (lower). Regions along the caecum are annotated and numbered; region 2, used in subsequent analyses, is indicated by a white rectangle. Scale bar, 1 mm. **i**, Changes in peristaltic contractions under the indicated pharmacological conditions: DMSO (1:1000), nifedipine (30 μM), or aphidicolin (10 µM). **j**, Frequencies of peristaltic waves in caecal fragments under the conditions shown in i. Data were analyzed using Welch’s t-test. **k**, Morphological changes in caecal fragments from E10 (upper) to E10 + 3 days in culture (lower) under the same conditions as in **j**. Scale bar, 500 µm. **l**, Growth rates (%) in length (left) and width (right) of caecal fragments under the same conditions as in **j** (mean ± SE).

We here used the caecum, a pair of gut tubes that protrude from the main tract and extend in straight (Fig. 1c) ^7,8^. At E8 to E12, the length of the caeca increases more rapidly than its width showing a typical pattern of anisotropic growth (Fig. 1d). The elongation rate is higher during E10–E12 (1.91-fold) than during E8–E10 (1.74-fold). Importantly, this accelerated increase in elongation at E10–E12 coincides with robust gut peristalsis (Fig. 1e, Movie 1, 2). These observations lead to an attractive hypothesis that peristaltic activity generated by the gut promotes its own elongation during embryogenesis. Using an organ culture combined with optogenetic technology we recently developed ^7^, we manipulated peristaltic activity during long-term culture, and found that peristalsis plays a critical role in the caecum elongation. Furthermore, live imaging of smooth muscle cells in the growing caecum reveals that peristaltic activity rearranges circumferentially proliferating smooth muscle cells to a longitudinal alignment. Thus, spontaneous muscle contractions in the form of peristaltic activity act as a growth signal to enable the anisotropic elongation of the embryonic gut.

## Materials and methods

### Chicken Embryos

Fertilized chicken eggs were purchased from the Yamagishi poultry farms (Wakayama, Japan), and embryos were staged using Hamburger and Hamilton (HH) series (Hamburger and Hamilton, 1951) or embryonic day (E). All animals were treated with the ethical approval of Kyoto University (#202508).

### Measurement of embryonic body anatomy

E10 and E12 embryos were imaged after being immersed in PBS for over 30 minutes to induce muscle relaxation. Images were taken using an iPhone and a Leica MZ10 F microscope equipped with a DS-Ri1 camera (Nikon) and analyzed using ImageJ (NIH). The length of body was measured from the tip of the beak to the tip of the tail along the midline of the back. The length of forelimb was measured from the elbow joint to the tip of the major digit (digit III) tracing the bones through the tissue. Eye was measured with its diameter. The length of gut was measured from esophagus to cloaca.

### Gut dissection

Three types of specimens were prepared: Intact caeca connected to the main tract was prepared by transecting at hindgut and from embryo with its mesentery removed; Whole caecum was prepared by cutting caecum at the junction between the caecum and the main tract; Caecal fragment was prepared by fragmenting caecum into 8 pieces with a similar length, and retrieving the middle fragment of the proximal part of the caecum, as mentioned as “P2” in Kawamura et al. 2025.

### Measurement of the dimensions of caecum

All gut specimens were immersed in phosphate buffered saline (PBS: 0.14 M NaCl, 2.7 mM KCl, 10 mM Na_2_HPO4–12H2O, 1.8 mM KH2PO4) more than 30-minutes before each measurement, in order to induce muscle relaxation. While measurement, the vertical curvature of the long caecum was gently straightened by lowering the PBS water level. The length of the intact gut connected to the main tract was measured from the branching point to the tip along the centerline. The width was calculated by dividing the area of the gut region by the length. The length of the whole caecum was measured from the amputated end and the tip along the centerline. The length of the P2 fragment was measured between two amputated ends along the centerline, and the width was measured at the central portion of its tubular structure.

### Organ culture and time-lapse imaging of gut contraction

For organ culture, intact gut connected to the main tract or whole caecum was placed within PDMS canals (internal dimensions: 0.8 mm x 1 mm) in a glass bottom Petri dish (3.5 cm diameter, IWAKI, 3960-035) to prevent drift, while P2 fragment was placed directly on a glass bottom. This specimen was cultured in 5 ml of DMEM/Ham’s F-12 with L-glutamine and Sodium Pyruvate (Wako, 045-30665) supplemented with 1× Penicillin-Streptomycin-Amphotericin B Suspension (PSA) (FUJIFILM, 161-23181). The dish was maintained in a heating chamber (BLAST, C-140T and BE-051A) at 38.5℃ under an atmosphere of 5% CO_2_ and 95% O_2_. High concentration oxygen was supplied by an oxygen generator (KMC, M1O2S10L). When culturing P2 fragment, Nifedipine (30 µM for a final concentration, 100 mM stock in DMSO, Wako, 141-05783), Aphidicolin (5 µM for a final concentration, 5 mM stock in DMSO, Cell Signaling, 32774S) or Ani9 (50 µM for a final concentration, 10 mM stock in DMSO, Sigma, SML1813-5MG) was administered, while DMSO (Wako, 037-24053) was added at 1/1000 (v/v) as a control. During culture, the P2 fragments were gently rolled with forceps once daily to prevent the serosa from adhering to the glass bottom. During culture, time-lapse images were acquired from above every 1-second using a Leica MZ10 F microscope equipped with a DS-Ri1 camera (Nikon). Because of the photodegradability of Nifedipine, time-lapse imaging was performed under low-light conditions for all analyses of the fragment.

### Motion analysis of gut

Time-lapse videos of intact gut connected to the main tract or whole caecum were analyzed with ImageJ to make kymographs, tracing from junctional point to the tip of the caecum. The number of peristalses in the caecum was counted for 1 hour at the middle region of the proximal part, as mentioned as P2 region in Kawamura et al. 2025. In contrast, in P2 fragment, peristalses were quantified by analyzing the change in the brightness of the P2 fragment region in movies. The signal polarity was adjusted such that spikes pointed upward, because whether the region became brighter or darker during contraction was arbitrary. For 3-days culture, the number of peristaltic events was counted over a one-hour window in the middle of each day (12hr-13hr、36hr〜37hr, and 60hr-61hr), and the values from the three days were averaged. For 24h culture (Fig. 2), the number of peristaltic events were detected and counted using SciPy library (scipy.signal. find_ peaks) in Python (parameters: height = 1.5).

**Figure 2:**
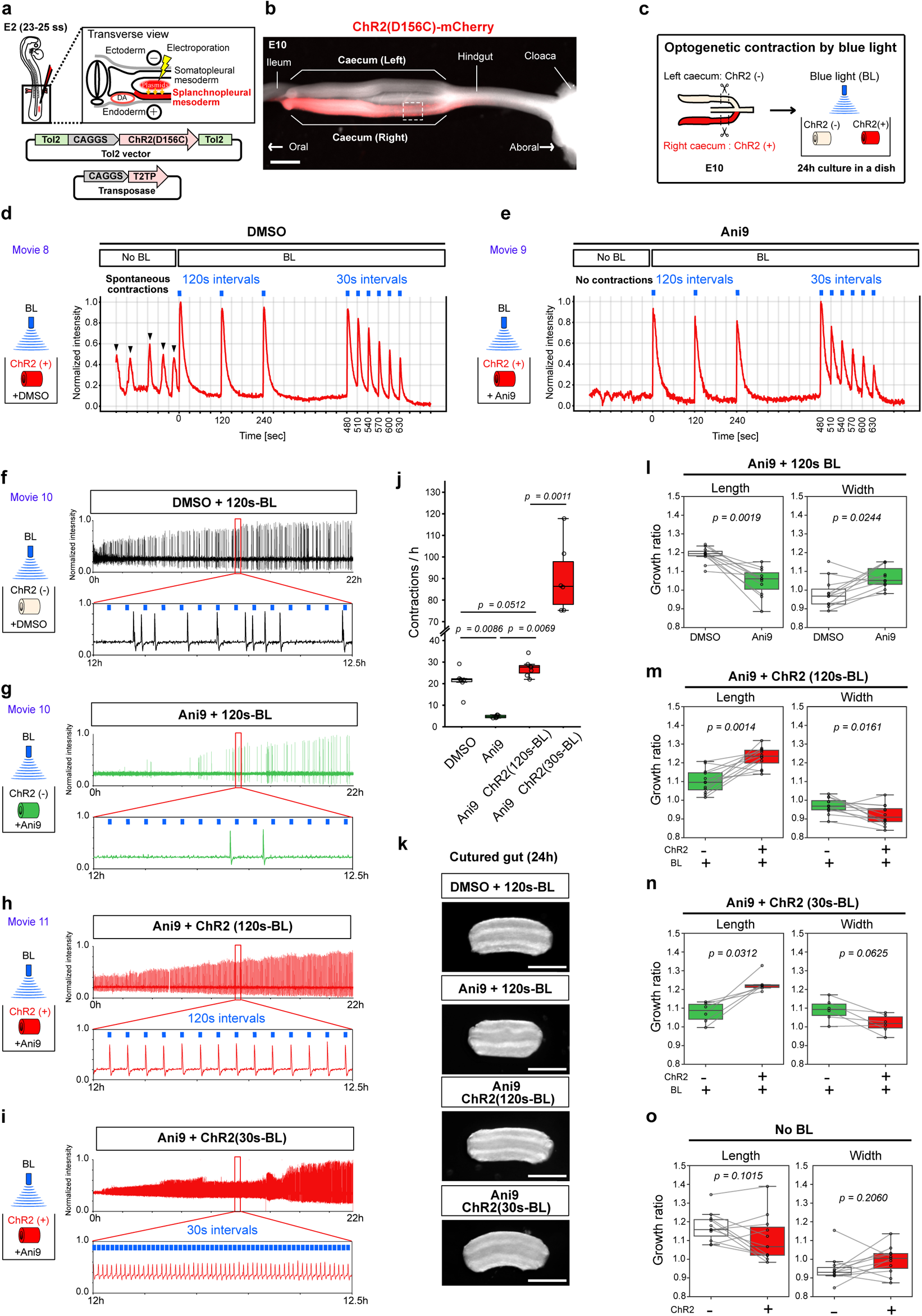
Optogenetically-triggered peristaltic contractions are sufficient for caecum elongation. **a**, Genetic introduction of ChR2 into caecum by in-ovo electroporation. **b**, Expression of ChR2(D156C)-mCherry in the right caecum. Scale bar, 1 mm. **c**, Blue-light stimulations of ex-vivo cultured caecal (P2) fragment with/without ChR2 (+/-). **d**, Optogenetic induction of peristaltic contractions in the control (DMSO, 1:200). The red trace indicates changes in fragment contraction. Black arrowheads mark spontaneous contractions, and blue bars indicate the timing of blue-light illumination. **e**, Optogenetic induction of peristaltic contractions in the presence of Ani9 (50 µM). Note that spontaneous contractions are suppressed, whereas responsiveness to blue light is retained. **f-i**, Long-term (22h) responses of ChR2(-) or ChR2(+) fragments to blue-light pulses under the indicated conditions: **f**, ChR2(-), DMSO (1:200), 120-s interval; **g**, ChR2(-), Ani9 (50 μM), 120-s interval; **h**, ChR2(+), Ani9 (50 μM), 120-s interval; **i**, ChR2(+), Ani9 (50 μM), 30-s interval. **j**, Frequencies of the contractions under each condition: DMSO, n = 6; Ani9, n = 6; Ani9 ChR2 (120s-BL), n =7; Ani9 ChR2 (30s-BL), n =6. **k**, P2 fragments after 24 h of incubation under the indicated conditions. Scale bar, 500 μm. **l-o**, Growth ratio (length and width) after 24h of incubations under the indicated conditions: **l**, n=11; **m**, n =12; **n**, n = 6; **o**, n = 12. Boxplots show the median (center line), IQR (25th–75th percentiles), and whiskers indicate 1.5xIQR. Statistical tests: **j**, Kruskal-Wallis test followed by Dunn’s multiple-comparison test with Holm correction; **l-m**, two-side Wilcoxon signed-rank test.

### In ovo electroporation

In ovo electroporation was performed as previously reported (Momose et al., 1999; Atsuta et al., 2013) with slight modifications (Shikaya el al. 2023, Kawamura et al. 2025). Briefly, a DNA solution was prepared at a concentration of ∼10 μg/μL, with a Tol2 plasmid : pCAGGS-T2TP: 4% fast green FCF (Wako, CI 42053) at a ratio of 4:1:0.5. Tol2 plasmids used in this study: pT2A-CAGGS-ChR2(D156C)-mCherry-IRES-Neor, pT2A-CAGGS-GAP-Orange-2A-H2B-EGFP. DNA solution was injected into the coelomic cavity of the lateral plate mesoderm on the right side of Hensen’s node (presumptive right caecum) of E2.5 (HH13-15) stage embryos (Shikaya et al., 2023). An electrical pulse of 50 V, 0.05 ms duration was applied, followed by five pulses of 7 V, 25 ms duration, at 250 ms intervals (BEX, Pulse generator CUY21EDIT II). Electroporated embryos were raised in an incubator at 38.5℃ with high humidity until the E10 stage.

### Optogenetic manipulation of peristalsis

A pair of caeca, with the right caecum expressing ChR2(D156C)–mCherry, was obtained from E10 embryos electroporated as described above. The right caecum expressing ChR2(D156C) served as the experimental group, and the left caecum served as an internal control. Both caeca were dissected from the main tract, and the P2 fragments were used. P2 fragments were subjected to the organ culture described above. Blue light (λ = 470 nm) was generated by an LED (OSB56L5111P) and delivered to the P2 fragment cultured in a glass-bottom dish. Trains of light pulses (100-ms pulse width, 5 Hz, 2-s train duration) were applied at 120- or 30-s intervals, controlled by a microcomputer (Arduino Uno; arduino.cc). The light power density was approximately 400 μW/mm^2^, which was measured by a power meter (Thorlabs, PM160). All optogenetic experiments were conducted under the red light (λ = 590 nm, Opto Code, EX-590 and LED-EXTA).

### Whole mount immunostaining

After muscle relaxation and 4% paraformaldehyde (PFA) fixation for 1 day at 4℃, the specimen was washed with 0.25% PBST for 15 minutes, twice and blocked with 0.05% blocking reagent overnight at 4℃. Then the specimen was incubated overnight at 4℃ with dilution of 1:400 anti-αSMA (Sigma Aldrich, A5228). Following washing in 0.25% PBST for 1 hour, twice at 4℃, the specimen was incubated overnight at 4℃ with 1:400 species-specific secondary antibody and 1:1000 DAPI. After washing in 0.25% PBST for 1 hour, three times at 4℃, the tubular specimen was opened using a tungsten wire (Nilaco, 461167, 0.10 mm diameter). The opened specimen was placed on the glass bottom of the dish with the serosa side facing downward, and gently pressed against the glass bottom with a silicon plate weighing 1.5 g. 2D Fluorescence images were acquired using A1 (Nikon).

### Section immunostaining

P2 specimen freshly dissected or after organ culture was immersed in PBS for over 30 minutes to induce muscle relaxation. After that, the specimen was fixed in 4% PFA at 4℃ overnight, washed with PBS, and then dehydrated in 15% sucrose/PBS overnight at 4℃. The sucrose solution was then replaced with OCT compound (Leica biosystems, 3801480), mounted into blocks, and transverse cryosections were obtained at 10 µm. Section sample mounted on a glass slide was washed and permeabilized in 0.25% Triton X (AlfaAesar, A16046) in PBS (PBST) for 5 min, three times at RT, followed by blocking with 0.05% blocking reagent for 15 min at RT. Then the specimen was incubated overnight at 4℃ with dilution of 1:200 anti-αSMA (Abcam, ab5694), and optionally with 1:400 anti-pHH3 Ser10 (Proteintech, 66863-1-1g). Following three times washing in 0.25% PBST for 5 min at RT, the specimen was incubated for 2 hours at RT with 1:400 species-specific secondary antibodies and 1:1000 DAPI (nacalai tesque, 11034-56). After washing three times in 0.25% PBST for 10 min at RT and mounting with Fluoromount (Diagnostic Biosystems, K024), 2D fluorescence image was acquired using BZ-X810 Fluorescence Microscope (Keyence) and Nikon A1R confocal microscope (Nikon). To maintain consistent color usage, green and red color in some images were flipped digitally so that smooth muscle tissue is visualized in red.

### Measurement of muscular cross-sectional area, cell concentration, and frequency of mitosis

The cross-sectional area of the muscular layer was measured from the αSMA-stained region of caecal sections. Cell concentration was measured in both transverse images from section staining and lateral images from whole-mount staining, placing a 20µm × 20µm square ROI within the muscular layer. DAPI signals within the ROI were counted according to the following criteria: (1) signal regions with and area ≥ 10 µm^2^ were counted; (2) for each region, if less than 30% of its whole area placed within the ROI, it was counted as 0 nucleus; if 30-70% placed within the ROI, it was counted as 0.5 nucleus; and if ≥70% placed within the ROI, it was counted as 1 nucleus. The frequency of cells under mitosis was measured in transverse images, by dividing the number of pHH3-DAPI double positive signals observed within the smooth muscle layer by the area of it.

### Cell morphology analysis

The P2 region expressing GAPOrange/ H2B-EGFP was prepared at E10, and muscle relaxation in PBS was introduced just after dissection or after three days of culture. Relaxed P2 was opened using a tungsten wire, and fixed in 4% PFA at 4℃ overnight. Following washing in PBS for 1 hour, the opened specimen was placed on the glass bottom with the serosa side facing downward, and pressed with a silicon plate. 3D Fluorescence images were acquired using a Nikon A1R confocal microscope, with z-stacks collected at 2 µm intervals. Spindle-shaped cells with identifiable poles were selected for measurement of cell minor axes and major axes. The minor axis was measured perpendicular to the midline of the thickest part of the cell, even if the cell was curved. The major axis was defined as the straight-line distance between the two poles, even if the z-axis (radial) positions of the poles differed. Cells for which only one axis could be measured were included only for that axis.

### Time-lapse imaging of cellular dynamics

GAP-Orange / H2B-EGFP expressing P2 fragments dissected from E10 embryo were placed on a glass bottom of 3.5 cm Petri dish, containing 5 ml of DMEM with DMSO (1/1000, v/v), Nifedipine (30 µM), or Aphidicolin (5 µM). Two silicone blocks secured to the bottom were placed on both lateral sides of the specimen holding it gently to prevent drift when the stage was moved. The specimen was maintained in a heating chamber at 38.5℃ under an atmosphere of 5% CO_2_ and 95% O_2_, and fluorescence time-lapse images were acquired from the bottom every 10-minutes using BZ-X810 Fluorescence Microscope (Keyence).

### Cell tracking and analysis

Cell positions were quantified based on the centroid of the nucleus. Circumferential positions were defined as the radial distance from the gut centerline, and longitudinal positions were defined relative to the positions of multiple cells along the gut axis. In one case where the P2 fragment rolled during culture (1 of 4 samples in the DMSO condition), the sample was excluded from circumferential analyses (Fig. 4) but retained for longitudinal analyses (Fig. 5). Nuclei of smooth muscle cells were identified by their spindle-shaped morphology labeled with GAP-Orange, with reference to images from neighboring time points or adjacent z-planes as needed, because some interstitial cells, in addition to smooth muscle cells, were fluorescently labeled. When image acquisition coincided with gut contractions and produced motion-blurred frames, particularly in DMSO-treated samples, images acquired within ±20 min of the intended time point were used instead.

Circumferential migration of smooth muscle cells was analyzed by generating circumferential kymographs over 5 h, using cell positions sampled every 20 min. For each cell, the relative position of the nuclear centroid was expressed as a percentage of the distance between the two cell poles (set to 100%). Circumferential migration velocity was calculated by converting the total circumferential displacement over 6 h into a rate per hour.

To analyze cell division orientation, the angle between the longitudinal axis of the gut and the line connecting the centers of the two daughter nuclei at the time of nuclear separation was measured. For daughter-cell tracking along the longitudinal axis, the distance between the two daughter cells was recorded at 30 min and 6 h after mitosis.

### Divergence index (DI)

Five smooth muscle cells that did not undergo mitosis during the imaging period were selected within a rectangular ROI (50 µm in width) along the longitudinal axis and tracked for 12 hours. The variance of their longitudinal positions at each time point was defined as follows: (i) the longitudinal coordinates of individual cells were extracted; (ii) the mean position of these cells (centroid) was calculated; and (iii) the mean squared difference from the centroid at time T was defined as V_t_. The divergence index (DI) was then defined as the natural logarithm of the ratio of variances at 12 hours and 0 hours, DI = log (V₁₂ / V_0_).

### Statistical analysis

Graphs were made by Excel or matplotlib and seaborn library in Python. Kruskal-Wallis test followed by Dunn’s multiple-comparison test with Holm correction, two-side Wilcoxon signed-rank test and Mann–Whitney U test were performed by library in Python.

## Results

### Inhibition of peristaltic movements distorted caecum elongation

A caecum prepared from E10 embryo was organ-cultured for 3 days. Peristaltic movements were retained and the caecum exhibited significant elongation of 1.56-fold. (Fig. 1f, g, Movie 3, 4). For quantitative assessments with high resolution, we used a short fragment “P2”, the second most proximal fragment among 8 chopped pieces ^8^ (Fig. 1h). P2 was organ-cultured for 3 days with each of following chemical inhibitors: nifedipine (a L-type Ca²⁺ channel inhibitor) to arrest peristalsis ^10,11^, aphidicolin (a DNA polymerase inhibitor) to arrest cell proliferation, and DMSO as a control. In control and aphidicolin groups, active peristaltic contractions were detected, whereas peristalsis was completely abolished in the nifedipine-treated P2 (Fig. 1i, j, Movie 5, 6, 7). We quantified changes in P2 morphology by measuring the length and width before and after the culture (Fig. 1k, l). Control P2 elongated by 1.44-fold with little changes in width. In contrast, nifedipine-treated P2 showed limited elongation (∼1.12-fold) accompanied by a significant increase in width (∼1.18-fold), which is consistent with the previous study with the midgut in chickens ^11^. Notably, despite robust peristaltic activity, aphidicolin-treated P2 exhibited modest elongation (∼1.15-fold) with a reduction in width (∼0.85-fold) (Fig. 1k, l). Thus, it is likely that peristaltic contractions are required for efficient longitudinal growth, whereas overall tissue growth depends on cell proliferation.

### Optogenetically induced peristaltic movements are sufficient to drive the gut anisotropic elongation

The possibility cannot be excluded that nifedipine affected not only peristalsis but also other unknown signals that are relevant to the P2 elongation. To unambiguously clarify the role of peristalsis in gut elongation, we experimentally induced peristaltic movements in P2 by optogenetic technology that we recently developed (Shikaya et al. 2023, Kawamura et al, 2025). For the optogenetic manipulation of gut movement in chicken embryonic gut, a modified type of channelrhodopsin 2, ChR2(D156C), serves as a powerful tool ^7,8,12^. Expression vector harboring ChR2(D156C) was unilaterally electroporated into splanchnopleural mesoderm (progenitors of gut mesoderm) of early chicken embryos (E2) (Fig. 2a, b). P2 fragment prepared from ChR2(D156C)-expressing E10 caecum [ChR2(+)] was subjected to optogenetic manipulations under blue-light illumination (Fig. 2c), whereas non-electroporated contralateral caecum [ChR2(-)] served as an internal control within the same embryo. This is critical because both the spontaneous peristaltic frequency and the extent of gut elongation vary between individuals.

Without blue light illumination, ChR2 (+) P2 exhibited the spontaneous contractions (Fig. 2d, No BL, and Movie 8) similar to those observed in Fig. 1i, j. When blue light pulses at 120-s intervals were delivered to the same specimen, each pulse reliably triggered a peristaltic contraction (Fig. 2d, BL and Movie 8). Such responses also occurred with 30-s intervals. To obtain P2 that implements solely optogenetically evoked contractions without spontaneous ones, we attempted to test a nifedipine-treated P2 (Fig.1i, j) for the optogenetic manipulations. However, blue lights failed to evoke any contractions probably because L-type Ca^2+^ channel (a target of nifedipine) is a downstream effecter of ChR2. We alternatively used Ani9, an inhibitor for Ca^2+^ -activated Cl^-^ channel (Ano1/TMEM16A), known to attenuate peristalsis ^13,14^. Addition of Ani9 to the culture medium drastically suppressed spontaneous peristaltic contractions of ChR2 (+) P2 (Fig. 2e, No BL and Movie 9). Markedly, with blue light pulses, this specimen elicited robust light-induced peristaltic contractions in a way similar to the control (Fig. 2e and Movie 9).

ChR2 (-) P2 fragments were organ-cultured for 24 h with blue light pulses at 120-s intervals. Frequencies of peristalsis in (DMSO (control)- and Ani9-P2 fragments were 20–30 and 5 events per hour, respectively (Fig. 2f, g, j and Movie 10). When the spontaneous peristalsis-free (+Ani9) ChR2 (+) P2 was stimulated with blue light with 120-s intervals, this fragment successfully responded to the light pulses at a frequency of ∼30 events per hour (Fig. 2h, j and Movie 11) comparable to the ChR (-) (DMSO) (Fg.2f, j), and ∼90 events per hour with 30-s intervals(Fig.2i, j).

We quantified changes in growth of these optogenetically treated P2 fragments after 24 h of organ culture. As already shown in Fig.1, control P2 [ChR2(+), DMSO] elongated well with little changes in width (Fig. 2k, l, DMSO). Ani9-treated ChR2 (-) P2 showed little elongation with increased width (Fig. 2k, l, Ani9), showing that suppression of peristalsis by Ani9 inhibits elongation, which is consistent with results of nifedipine-treated experiments (Fig. 1k, l). Strikingly, when peristalsis was re-activated in Ani9-treated fragments by 120-s interval light stimulation, the elongation rate was restored to the level comparable to the control (length ∼1.2-fold, width ∼0.9-fold; Fig. 2k, m). Light stimulation at 30-s intervals yielded a similar degree of elongation (Fig. 2k, n). ChR2 expression gave no impact on elongation without light stimulation (Fig. 2k, o). Together, peristaltic contractions are sufficient to drive the gut anisotropic growth.

### Neither cell hypertrophy nor cell density changes account for the anisotropic growth of P2

To know how the peristaltic contractions drive anisotropic growth of the caecum, we investigated possible changes in circular smooth muscle (CSM) cells (Fig. 3a), since these cells are the primary target of optogenetic control (the outer most SM cells remain unaligned longitudinally at E10). As shown in Fig. 3b, a ring of CMS layer seen in transverse sections stained with αSMA is enlarged in nifedipine-P2, and smaller in aphidicolin specimens, mirroring the macroscopic observations shown in Fig. 1k and l. We subsequently compared the volumetric growth rates of the CSM layer between control and nifedipine-treated P2 fragments. To do so, a change rate of the cross-sectional area of the CSM layer after 3-day culture (E10 + 3) was multiplied by that of the longitudinal elongation (Fig. 1l). Notably, the volumetric growth rates were comparable between DMSO control P2 (1.26-fold) and nifedipine-treated one (1.30-fold). Thus, the volumetric increase of the CSM layer is independent of peristaltic activity. In contrast, aphidicolin treatment failed to increase in volume (0.95-fold), showing cell proliferation is needed for the gut growth as expected.

**Figure 3:**
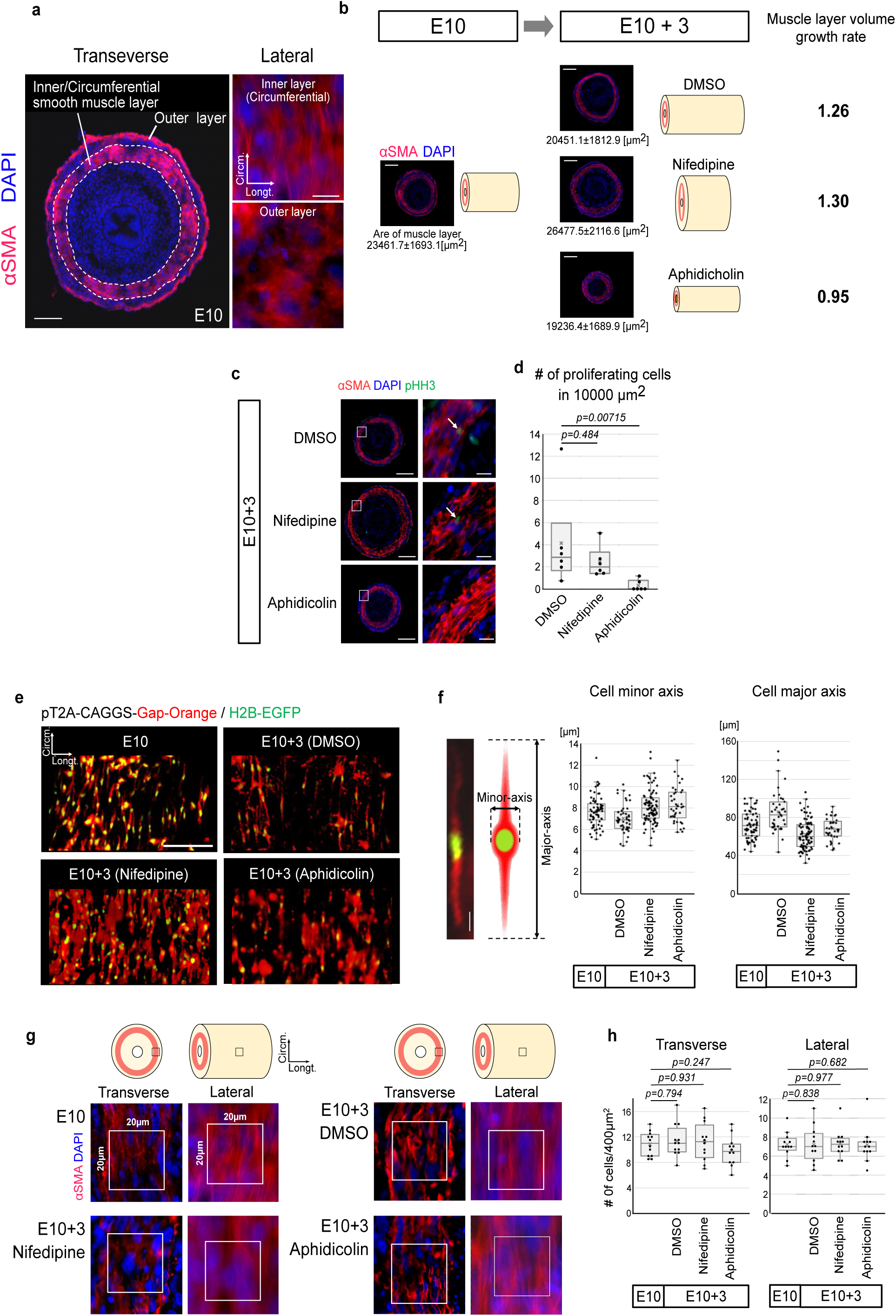
Neither cell hypertrophy nor cell density changes account for the anisotropic growth of caecum. **a**, Transverse cross-section and lateral views of whole-mount immunostaining of smooth muscle (aSMA, red) and nuclei (DAPI, blue) in E10 caecum. Dotted outlines display circumferential smooth muscle (CSM) layer. Scale bar, 50µm (Left panel); 10µm (right panels). **b**, Transverse cross-section of caecal (P2) fragments before (E10) and after (E10+3) cultured with DMSO, Nifedipine, or Aphidicolin. Red, α-SMA; blue, DAPI. Mean areas of CSM layer are shown below each image (n=4). Volume increase ratios are defined as (the cross-sectional area increase ratio) × (the length increase ratio). Scale bars, 100μm. **c**, Immunofluorescence staining of E10 + 3-day cultured P2 fragments for cell proliferation (pHH3, green), smooth muscle (αSMA, red), and nuclei (DAPI, blue). Inset, magnified images. White arrows show proliferating cells. Scale bars, 100 μm (left columns), and 10µm (right columns). **d**, Box plots of proliferating cell density [/10,000 µm²] under each condition (n = 6 for each). Statistical significance was assessed by Mann–Whitney U test (p < 0.05). **e**, Genetic labeling of CSM cells by in-ovo electroporation in P2 fragment. Fluorescent signals (red, Gap-Orange; green H2B-EGFP) are detected from lateral view of P2 fragment. E10 and E10+3 cultured fragment with DMSO, Nifedipine, or Aphidicolin are shown. Scale bar, 100 μm. **f**, The size of CSM cells before and after culture with drugs. Representative image of a smooth muscle cell labelled fluorescently (left). Schematic showing the definition of minor/major axis of a cell (middle), and box plots of cell minor/major axis length in the condition of E10 and E10+3 with DMSO, Nifedipine, and Aphidicolin (right two). In minor axis, n = 70 cells in total from 3 individuals (E10), n=49 cells in total from 3 individuals (DMSO), n=91 cells in total from 3 individuals (Nifedipine), and n=38 cells from 3 individuals (Aphidicolin). In the major axis, n=68 cells in total from 3 individuals (E10), n = 40 cells from 3 individuals (DMSO), n = 96 cells from 3 individuals (Nifedipine), and n = 38 cells from 3 individuals (Aphidicolin). **g**, The cell density of cells in circumferential smooth muscle layers. Representative images of E10, DMSO, Nifedipine, or Aphidicolin conditions, transverse and lateral view of caecum, stained against α-SMA (red) and DAPI (blue). The white squares (20 µm per side) indicate ROIs. Schematic illustrating transverse and lateral views are shown upper. **h**, Box plots of cell density [/400 µm²] in transverse (left) and lateral (right) views in the four conditions. n=12 cells from 3 individuals for all conditions. Statistical significance was assessed by Mann–Whitney U test.

We examined cell proliferation by phospho-histone H3 (pHH3) used in growing gut of chicken embryos ^15^ (Fig. 3c, d). Distribution of pHH3-positive cells was comparable between control (DMSO) and nifedipine-treated P2, and as expected, abolished by aphidicolin (Fig. 3c, d). Other possibilities accounting for the anisotropic growth include anisotropic hypertrophy of individual CSM cells, and changes in their cell density. A CSM cell is spindle-like in shape, and its major and minor axes were measured for GAP-Orange/H2B-EGFP-electroporated cells. These two values did not account for the changes seen for the 3-day cultures in control-, nifedipine-, and aphidicolin-P2 fragments (Fig. 3, e. f). For cell density, αSMA- and DAPI-positive cells were counted in a given area both in transverse and lateral views (Fig. 3, g, h), yielding no detectable changes after 3-day culture in all the drug-treated P2 fragments. Thus, the anisotropic growth of P2 cannot be accounted for by cell hypertrophy or changes in cell density.

### CSM cells undergo circumferentially oriented cell division independently of peristaltic movements

To find out cellular dynamics that may account for the anisotropic growth of the gut, we performed live imaging of fluorescently labeled CSM cells (GAP-Orange/H2B-EGFP-electroporated) in cultured P2. To our surprise, CSM cells are actively translocated along the circumferential axis with some cells moving either direction (Fig. 4a, Movie 12). Kymography of nuclear position and cell shape indicates that this motility reflects movement of entire cells and not of intracellular nuclear displacement (Fig. 4a), known for interkinetic nuclear migration (elevator migration) in developing neuroepithelium. The circumferential movements of and its velocity of CSM cells were comparable between control and nifedipine-treated P2 (Fig. 4b, c, Movie 13).

**Figure 4:**
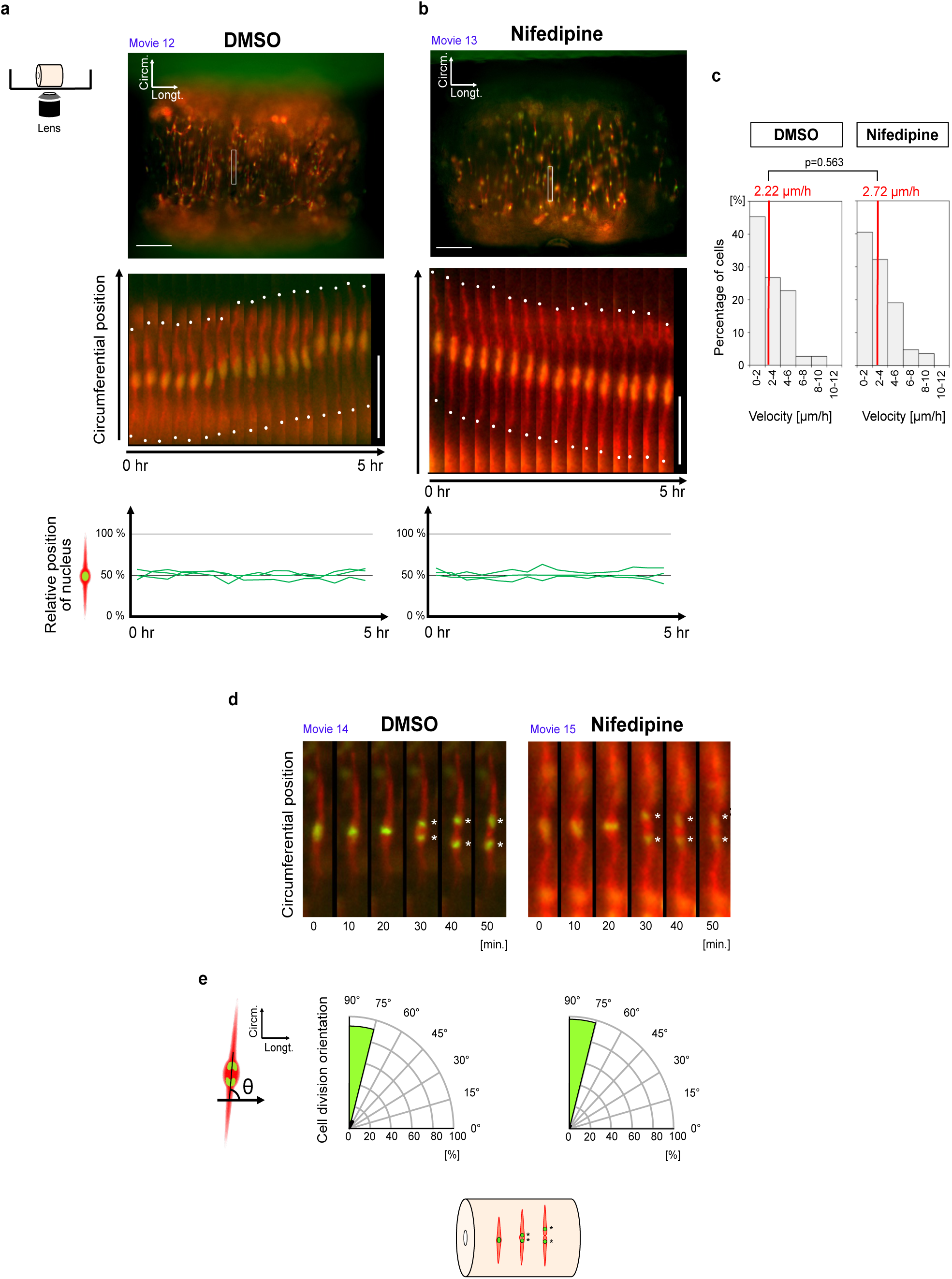
Peristalsis-independent circumferential migration and division of CSM cells. **a**, Representative image of P2 fragment labelled fluorescently (red, Gap-Orange; green H2B-EGFP) from lateral view (top panel), with a white box indicating a single cell selected for kymograph analysis. Scale bar, 100 µm. Kymograph profiles the movement of a smooth muscle cell along the circumferential axis with white dots on the cell poles (middle). Scale bar, 50 µm. The line graph shows the relative positions of nuclei within cells over time (lower). A schematic of the live-imaging system is shown left. (B) The same analyses as **b**, were conducted in Nifedipine condition. **c**, Histograms show the circumferential velocities of smooth muscle cells in DMSO (left) and Nifedipine (right) conditions. Red vertical lines represent medians, which were assessed statistically by Mann–Whitney U test. **d**, Kymographs showing a dividing smooth muscle cell in DMSO (left) and Nifedipine (right) conditions along the circumferential axis. The asterisks indicate the nuclei of newly generated cells. **e**, Quarter rose plots showing cell division orientation in DMSO (left) and Nifedipine (right) conditions. Schematic defining cell division orientation is shown left, and graphical abstract of smooth muscle cell mitosis is shown below.

CSM cells undergo mitosis. Notably, our live imaging analyses revealed that cells divided in a circumferential direction, which is perpendicular to the gut axis, and this mitotic orientation was not affected by nifedipine-treatment (Fig. 4d, e, Movie 14, 15). This unprecedented cell division profile highlights that the proliferative input initially expands the CSM layer circumferentially, and raises the possibility that the CSM layer would keep growing circumferentially unless peristaltic movements take place, which agrees with the observation for the length and width shown in (Fig. 1k, l). Thus, we reasoned that the peristalsis might be required for the circumferentially proliferating CSM cells to be rearranged to the longitudinal alignment.

### Peristaltic contractions cause longitudinal rearrangement of CSM cell population

To test our hypothesis, we live-tracked CSM cells in cultured P2. We first tracked newly generated daughter cells to see whether they become separated along the longitudinal axis of the caecum. At 30 min and 6 h after mitosis, the two daughter cells were indeed separated longitudinally, but this profile was not affected by nifedipine-P2 (poorly elongating) (Fig. 5a).

**Figure 5:**
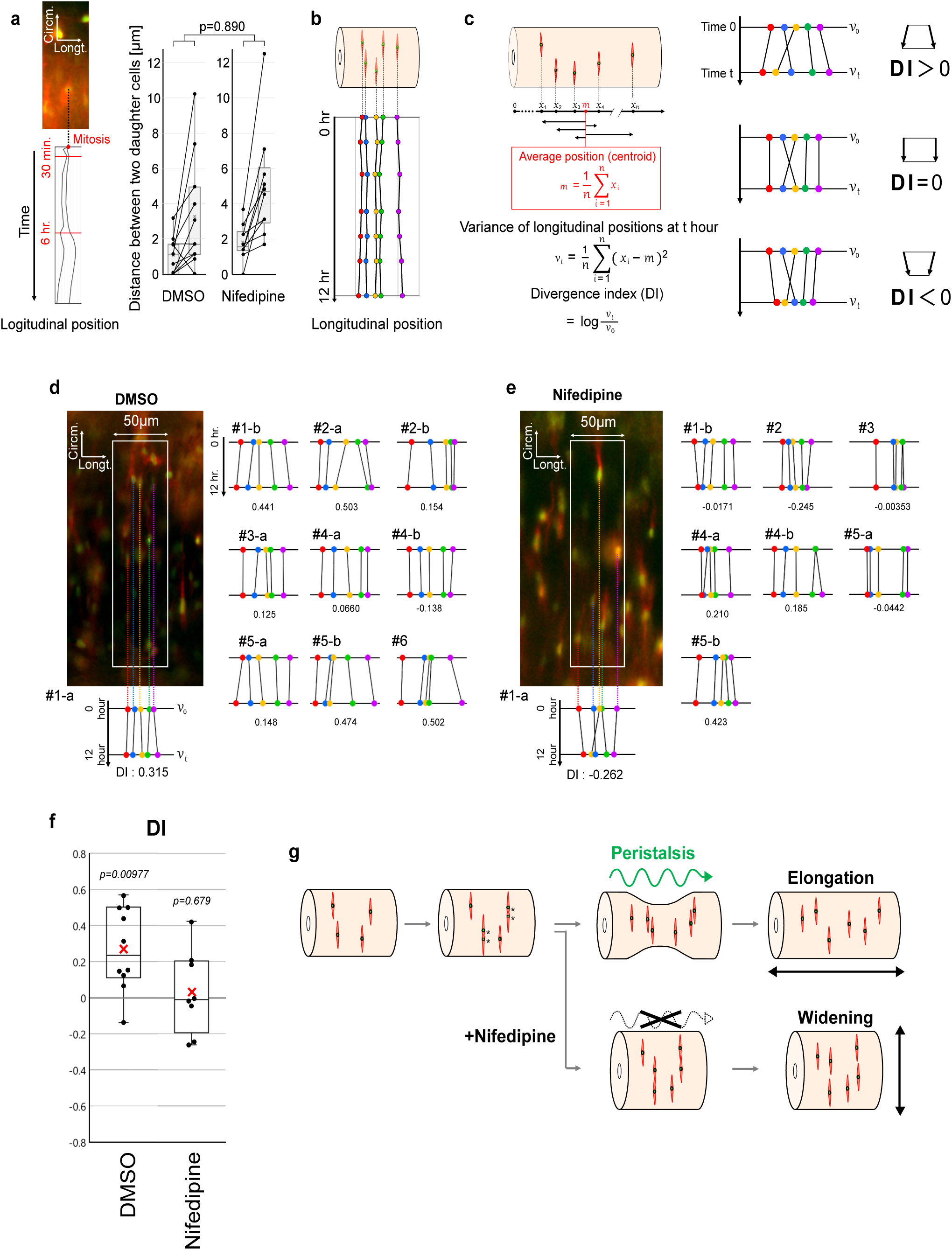
Peristalsis-dependent longitudinal rearrangement of CSM cell population. **a**, The daughter cell tracking. The longitudinal trajectories of daughter cells after mitosis, with the image of a cell at the verge of mitosis shown above (left). Time points 30 minutes and 6 hours after mitosis are indicated with red lines. Paired scattered plots overlaid on box plots show the profiles of distances between daughter cells 30 min. and 6 hr. after mitosis in DMSO (left) and Nifedipine (right) conditions (right). **b**, The longitudinal trajectories of a cell population. **c**, Divergence analysis. Definition of Divergence Index (DI) is shown left, while the meaning of DI is summarized in the right column. **d**, The divergence analysis in DMSO condition. Representative image of the fluorescently labeled P2 fragment from the lateral view, with white rectangle indicating the region for cell sampling, and with the trajectories from time 0 hr. to time 12 hr. and DI shown below (left column). Trajectories and DI in other samples (right). **e**, The divergence analysis in Nifedipine condition. **f**, Box plots of DI. **g**, Summary figure illustrating how the peristalses contribute to the circumferential elongation of the gut, and what changes arise upon inhibition of peristalsis. Statistical tests: **a**, Mann–Whitney U test comparing distance changes between two conditions; f, one-sample Wilcoxon signed-rank test against 1.00.

We next assessed collective changes/rearrangement within a broader CSM cell population rather than immediate post-mitotic daughter cells (Fig. 5b). For this purpose, we introduced an index reflecting redistribution, the Divergence Index (DI), defined as the log change in variance of longitudinal cell positions between the start and end of the analysis (Fig. 5c) A positive DI (> 0) indicates a divergence of cell positions along the longitudinal axis, whereas a DI near zero (= 0) means little net redistribution. A negative DI (< 0) is of longitudinal convergence of cell position.

Using this metric, we live-tracked five cells in each specimen within a 50 µm-wide region over 12 h. In control P2 fragment, DI was positive (median 0.234, p = 0.00977), indicating a longitudinal divergence of CSM cell positions (Fig. 5d, f). In contrast, nifedipine-treated P2 showed no significant change in DI (median 0.0103, p = 0.945) (Fig. 5e, f). These results indicate that peristaltic contractions promote longitudinal redistribution of CSM cell population, and that such tissue-scale cell rearrangements enable the gut anisotropic growth. Thus, embryonic peristaltic contractions are not merely a rehearsal for diet intake after hatch, but play a pivotal role in directing circumferentially dividing cells to be rearranged along the longitudinal axis leading to a massive elongation of this organ.

## Discussion

We have shown that the peristaltic movements direct anisotropic growth of the embryonic gut, which would otherwise keep expanding radially. The anisotropic growth is accounted for by unexpected cell dynamics: CSM cells divide in a direction perpendicular to the gut axis followed by a collective and divergent distribution of cells along the gut axis, the latter taking place in a peristalsis-dependent manner. We propose a “two-step model” to explain the gut elongation: the volume increase is achieved by circumferentially proliferating cells, and peristaltic movement, in turn, reorients the CSM layer in 90 degrees. Such mechanisms may provide an efficient way for the gut to grow longer while maintaining an appropriate diameter and functional integrity. It is plausible that divided CSM cells retain the circumferentially orientated spindle-like shape during the longitudinal rearrangement. In this way, the alignment of CSM cells along the circumferential axis is secure, which is critical to a local constriction during the gut tube peristalsis.

The CSM rearrangements demonstrated in this study bear conceptual similarity to convergent extension (CE) movements described in other embryonic tissues, such as cell intercalations in axial extension ^16^. In conventional CE, a tissue elongates with narrowing. We here cast a novel and parsimonious view that the massive elongation of the gut without jeopardizing the diameter of gut tube is underlain by the continuous supply of circumferentially oriented divisions prior to the collective cell rearrangement. Our findings give additional contribution to understanding shape changes, in particular, elongation of tissues and organs during body formation.

Each peristaltic cycle inevitably confers mechanical changes, such as cyclic strain and altered tissue tension ^5,10,11^, and their associated changes in intracellular signaling including Ca²⁺ dynamics ^17^. It has yet to be clarified how the mechanochemical coupling mediates the translation of peristaltic activity into changes in collective CSM rearrangement.

A possible contribution of other layers in the gut cannot be excluded: peristaltic stimulation by CSM might affect submucosa and endodermal epithelium contributing directly or indirectly to the massive elongation. And it is of interest to see if the two-step mechanism unveiled in this study is shared among other regions of the gut, for example, midgut and hindgut display distinct patterns of peristalsis ^9,18^.

## Acknowledgment

This work was supported by JSPS KAKENHI (Grant Numbers: 23H04702 for M. I. and 23H04933 for Y. T.), FY 2022 Kusunoki 125 of Kyoto University 125th Anniversary Fund for M. I. and Japan Science and Technology Agency (JST) FOREST Program (Grant Number: JPMJFR2334) for M.I.. K.K. is a fellow of JSPS (Grant Number: 25KJ1547). We thank Dr. Ishihara and Dr. Nakajima for helpful discussion, and Dr. Kawai for technical advice of chemical inhibitors. We also thank National Bio Resource Project (Chicken-Quail, Nagoya University) for their technical help.

## Author contributions

KK and MI conducted experiments. KK, MI, and YT wrote the paper. All authors listed have made a substantial, direct, and intellectual contribution to the work and approved it for publication.

## Movie legends

### Movie 1: Peristaltic movement in E9 caecum connected to the main tract, related to Fig. 1e

Time-lapse imaging of E9 chicken caecum connected to the main tract cultured ex-vivo, mounted in a PDMS cannel. Images were acquired every 1 second.

### Movie 2: Peristaltic movement in E11 caecum connected to the main tract, related to Fig. 1e

Time-lapse imaging of E11 chicken caecum connected to the main tract cultured ex-vivo, mounted in a PDMS cannel. Images were acquired every 1 second.

### Movie 3: Peristaltic movement in E10 whole caecum, related to Fig. 1f

Time-lapse imaging of E10 chicken whole caecum cultured ex-vivo, mounted in a PDMS cannel. Images were acquired every 1 second.

### Movie 4: Peristaltic movements in whole caecum cultured after 3 days from E10, related to Fig. 1f

Time-lapse imaging of chicken whole caecum after 3 days of ex vivo culture starting at E10, mounted in a PDMS cannel. Images were acquired every 1 second.

### Movie 5: Behavior of P2 fragment at E10 in DMSO, related to Fig. 1i

Time-lapse imaging of E10 chicken P2 fragments cultured with DMSO. Images were acquired every 1 second.

### Movie 6: Behavior of P2 fragment at E10 in nifedipine, related to Fig. 1i

Time-lapse imaging of E10 chicken P2 fragments cultured with nifedipine. Images were acquired every 1 second.

### Movie 7: Behavior of P2 fragment at E10 in aphidicolin, related to Fig. 1i

Time-lapse imaging of E10 chicken P2 fragments cultured with aphidicolin. Images were acquired every 1 second.

### Movie 8: Peristalsis in P2 fragment at E10 cultured with DMSO, illuminated with blue light, related to Fig. 2d

Channelrhodopsin-electroporated P2 fragments were cultured with DMSO. No blue light was applied during initial 3 minutes. Subsequent blue light stimulation was delivered three times at 120 s intervals, followed by six times stimulations at 30 s intervals. Images were acquired every 0.5 second.

### Movie 9: Peristalsis in P2 fragment at E10 cultured with Ani9, illuminated with blue light, related to Fig. 2e

Channelrhodopsin-electroporated P2 fragments were cultured with Ani9. No blue light was applied during initial 3 minutes. Subsequent blue light stimulation was delivered three times at 120 s intervals, followed by six times stimulations at 30 s intervals. Images were acquired every 0.5 second.

### Movie 10: Peristalsis of P2 fragment cultured with DMSO or Ani9, illuminated with blue light at 120-s intervals, related to Fig. 2f, g

Not-electroporated P2 fragments were cultured with DMSO or Ani9 (in separate dishes). Blue light was applied at 120-s intervals, images were acquired every 1 second.

### Movie 11: Peristalsis of P2 fragment expressing channelrhodopsin cultured with Ani9, illuminated with blue light at 120-s intervals, related to Fig. 2h

Channelrhodopsin-electroporated P2 fragments were cultured with Ani9 (in the same dishes). Blue light was applied at 120-s intervals, and images were acquired every 1 second.

### Movie 12: Cell dynamics in P2 fragment cultured from E10 in DMSO, related to Fig. 4a

GAP-Orange/H2B-EGFP-electroporated P2 fragment was cultured with DMSO. Fluorescence images were acquired every 10 minutes.

### Movie 13: Cell dynamics in P2 fragment cultured from E10 in nifedipine, related to Fig. 4b

GAP-Orange/H2B-EGFP-electroporated P2 fragment was cultured with nifedipine. Fluorescence images were acquired every 10 minutes.

### Movie 14: Mitosis of a smooth muscle cell in P2 fragment cultured from E10 in DMSO, related to Fig. 4d

GAP-Orange/H2B-EGFP-electroporated P2 fragment was cultured with DMSO. A dividing cell was magnified, and fluorescence images were acquired every 10 minutes.

### Movie 15: Mitosis of a smooth muscle cell in P2 fragment cultured from E10 in nifedipine, related to Fig. 4d

GAP-Orange/H2B-EGFP-electroporated P2 fragment was cultured with nifedipine. A dividing cell was magnified, and fluorescence images were acquired every 10 minutes.

